# Convergent evolution of ventral adaptations for enrollment in trilobites and extant euarthropods

**DOI:** 10.1101/2023.09.28.560020

**Authors:** Sarah R. Losso, Pauline Affatato, Karma Nanglu, Javier Ortega-Hernández

## Abstract

The ability to enroll for protection is an effective defensive strategy that has convergently evolved multiple times in disparate animal groups ranging from euarthropods to mammals. Enrollment is an evolutionary staple of trilobites, and their biomineralized dorsal exoskeleton offers a versatile substrate for the evolution of interlocking devices. However, it is unknown whether trilobites also featured ventral adaptations for enrolment. Here, we report ventral exoskeletal adaptations that facilitate enrollment in exceptionally preserved trilobites from the Upper Ordovician Walcott-Rust Quarry in New York State, USA. Walcott-Rust trilobites reveal the intricate three-dimensional organization of the non-biomineralized ventral anatomy preserved as calcite casts, including the spatial relationship between the articulated sternites (i.e., ventral exoskeletal plates) and the wedge-shaped protopodites. Enrollment in trilobites is achieved by ventrally dipping the anterior margin of the sternites during trunk flexure, facilitated by the presence of flexible membranes, and the close coupling of the wedge-shaped protopodites. Comparisons with the ventral morphology of extant glomerid millipedes and terrestrial isopods reveal similar mechanisms used for enrollment. The wedge-shaped protopodites of trilobites closely resemble the gnathobasic coxa/protopodite of extant horseshoe crabs. We propose that the trilobites’ wedge-shaped protopodite simultaneously facilitates tight enrollment and gnathobasic feeding with the trunk appendages.

## Introduction

The ability to completely enroll the body to form a tight protective ball to deter predatory attacks represents an effective strategy that has evolved multiple times throughout bilaterian evolution, including archetypical examples like xenarthan mammals (Superina and Loughry, 2012) and several euarthropods such as myriapods (Hannibal and Feldmann, 1981; Shear et al., 2011), terrestrial isopods (Brökeland et al., 2001; Hyžný and Dávid, 2017), and even some insect lineages (Ballerio and Grebennikov, 2016). Among extinct species, enrollment has been thoroughly documented in trilobites, a megadiverse group of marine euarthropods typified by the presence of a biomineralized calcitic dorsal exoskeleton (Fig. 1). Trilobite evolutionary history throughout the Paleozoic was heavily influenced by their ability to enroll effectively (e.g., Esteve et al., 2013; Suárez and Esteve, 2021; Chipman and Drage, 2023). Early Cambrian species show evidence of complete but imperfect (i.e. no encapsulating) enrollment, leaving open gaps between the thoracic and pygidial spines (Ortega-Hernández et al., 2013), whereas more derived groups evolved a diverse array of complex interlocking coaptive devices to make this defensive strategy more effective (e.g., Esteve et al., 2011, 2017, 2018). Despite its significance for the long-term evolutionary success of trilobites, enrollment is exclusively known from the perspective of the dorsal exoskeleton due to the paucity of enrolled specimens with exceptionally preserved non-biomineralizing ventral structures (Fig. 1). Thus, the precise physical mechanisms by which the ventral surface of trilobites could accommodate their numerous biramous appendages and other exoskeletal structures during enrollment remains enigmatic. Attempts to explain how the limbs would be organized relative to each other during enrolment have focused on hypothetical reconstructions of the non-biomineralized structures, like the exceptionally preserved Ordovician trilobite *Placoparia cambriensis* (e.g., Whittington, 1993). Although this reconstruction considered the position of the flexible intersegmental tendinous bars based on fossil data, it did not account for the presence of sternites (i.e., ventral exoskeletal plates) that are located in the medial space between each pair of limbs (Whittington, 1993). Moreover, trilobite appendages are infrequently preserved, being only known from *Konservat-Lagerstätten* such as the early Cambrian Chengjiang (e.g., Ramsköld and Edgecombe, 1996; Hou et al., 2008), mid-Cambrian Burgess Shale (e.g., Whittington, 1975; Losso and Ortega-Hernández, 2022), and Ordovician Beecher’s Bed (e.g., Whittington and Almond, 1987; Hou et al., 2021). Trilobite macrofossils from *Konservat-Lagerstätten* are typically highly compressed and their appendages are only found in prone specimens, limiting our understanding of the three-dimensional morphology and organization of the limbs during enrollment. In this context, the precise morphology of trilobite appendages has also not been comprehensively considered in terms of how they would fit in a completely enrolled position. The cross-section shape of the trilobite protopodite, for example, has been illustrated as either oval (Whittington, 1975; Whittington and Almond, 1987; Bruton and Haas, 1999; Bicknell et al., 2021; Schmidt et al., 2021) or square (Hou et al., 2021), or authors have omitted them altogether because of the lack of available data - (Whittington, 1993). While these differences may seem minor, shape plays a critical role in the function of various body parts (e.g., Bicknell et al. 2021), and thus this represents a fundamental gap of missing data when reconstructing the early autecology and functional morphology of one of the first successful clades in the evolutionary history of euarthropods.

**Figure 1.**
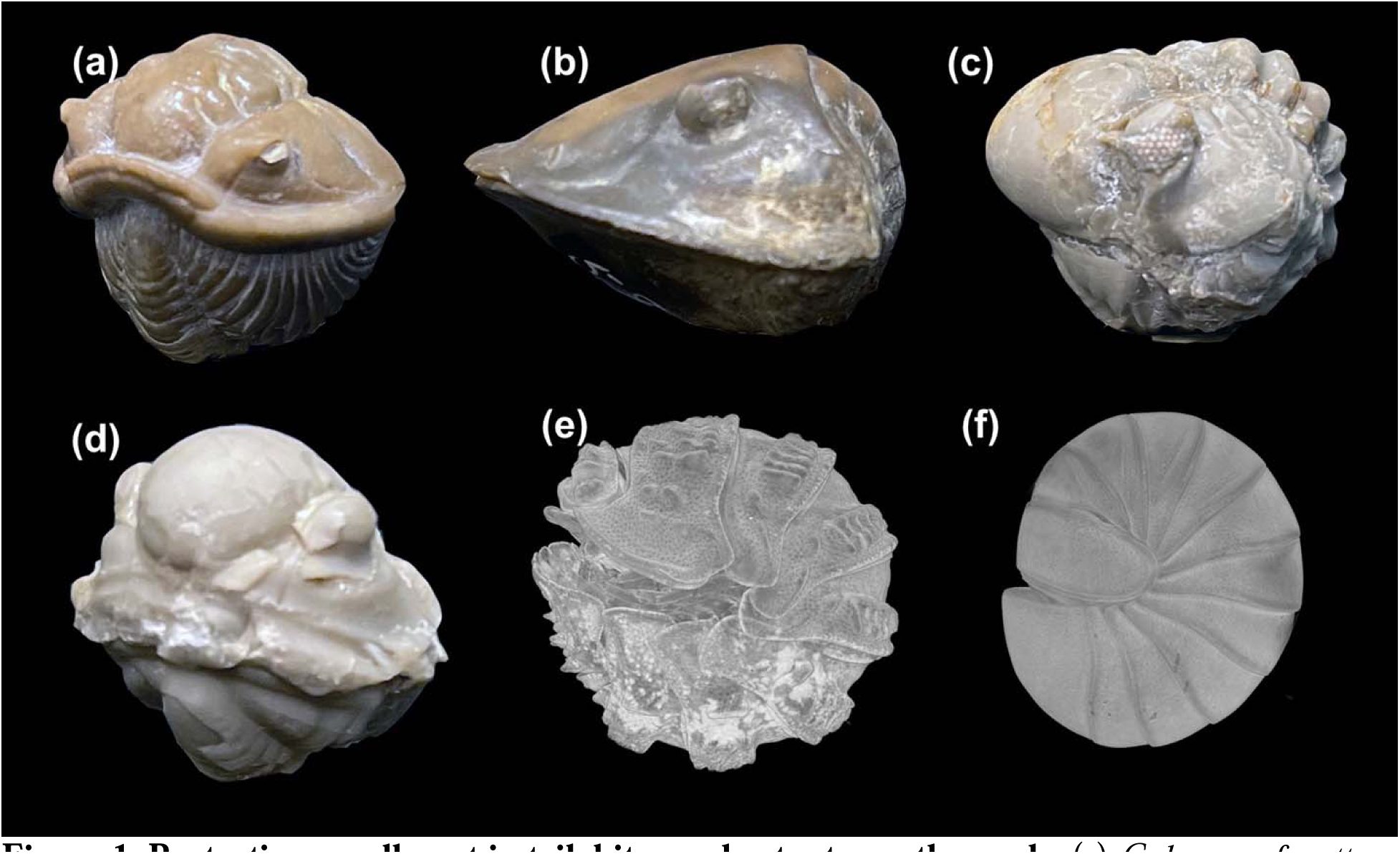
Dissection of terrestrial glomerid millipede. (a) Ventral view of complete specimen showing medial appendage attachment. (b) Ventral view of appendages. (c) Dissection of sternites and appendages with endopodites of right side removed. (d) Anterior view of dissected appendage pair with sternite and medial part of pleurite. Abbreviations: ple, pleurite; pt, protopodite; st, sternite.

In this study, we describe the non-biomineralized three-dimensional ventral exoskeletal morphology of trilobites from the Upper Ordovician (Mohawkian) Rust Formation of New York state based on exceptionally preserved fossils with soft tissues replicated as calcite casts (Losso et al., 2023). Trilobites from the Walcott-Rust Quarry are preserved in various stages of trunk flexure, revealing the intricate coupling of the biramous appendages and sternites to maximize a tight complete enrollment. Comparisons with the three-dimensional exoskeletal and appendicular morphology of extant euarthropods, including a glomerid millipede, a terrestrial isopod, and the Atlantic horseshoe crab *Limulus polyphemus*, indicate striking cases of convergent evolution in terms of their ventral exoskeletal anatomy. The presence of these functionally similar adaptations in phylogenetically disparate euarthropod lineages demonstrates a profound case of convergent evolution towards a common method of body enrollment separated by over 500 million years.

## Materials and methods

All studied specimens are housed at the Museum of Comparative Zoology (MCZ) at Harvard University (Cambridge, Massachusetts, USA). Trilobites from the Walcott-Rust Quarry originate from the Upper Ordovician Rust Formation, Trenton Group, in New York state. The exceptionally preserved trilobite fossils are composed of three-dimensional calcite casts of non-biomineralized tissues in a micritic limestone matrix from Layer 3 of the Rust Formation (see Losso et al., 2023). The studied trilobites are mounted as thin sections, produced by Charles D. Walcott in the 1870s (Yochelson, 1998). Fossil specimens were imaged at the Digital Imaging Facility (DIF) at the MCZ using a Keyence microscope with transmitted light. Extant euarthropod specimens sampled from the Invertebrate Zoology collections at the MCZ were imaged to analyze their protopodite and sternite morphologies, including *Limulus polyphemus* (MCZ:IZ:41112), a glomerid millipede (MCZ:IZ:165554) and a terrestrial isopod (MCZ:IZ:90105). All three specimens were stained with iodine prior to micro-computed tomographic scanning using a Bruker SkyScan 1173 micro-CT scanner at the DIF. Extant specimens were stained in iodine to increase resolution of Micro-CT scanning (See Supplemental Information for detailed staining method). Micro-CT imaging was performed at a voltage of 80 kV, wattage of 100 μA, a resolution of 6 μm, and with a 0.5 mm thick aluminum filter. Scans were reconstructed as TIFF stacks in NRecon (Bruker Corporation) and visualized and segmented in Dragonfly 2019 4.0 (Object Research Systems, Montreal, Canada).

One specimen of MCZ:IZ:165554 from the lot of six was dissected and photographed. Dissections were performed using an Ziess Stemi 305 microscope under direct light conditions, and photographs were taken using a Ziess Axiocam 208 color camera.

## Results

### Sternite morphology and preservation

Specimen MCZ:IP:158251, a thin section of the cheirurid trilobite *Ceraurus pleurexanthemus* in a completely enrolled position (Fig. 2a), reveals the three-dimensional morphology of ventral exoskeletal structures in exceptional detail. The presence of the hypostome and articulating half rings on the same specimen indicates that the section follows the sagittal plane along the midline of the body (Fig. 2a). In addition to showing the pattern of overlap and articulation of the tergites, MCZ:IP:158251 also preserves non-biomineralized ventral exoskeletal structures bound by the body wall as delimited by the presence of sparry and fibrous calcite (Losso et al., 2023).

**Figure 2.**
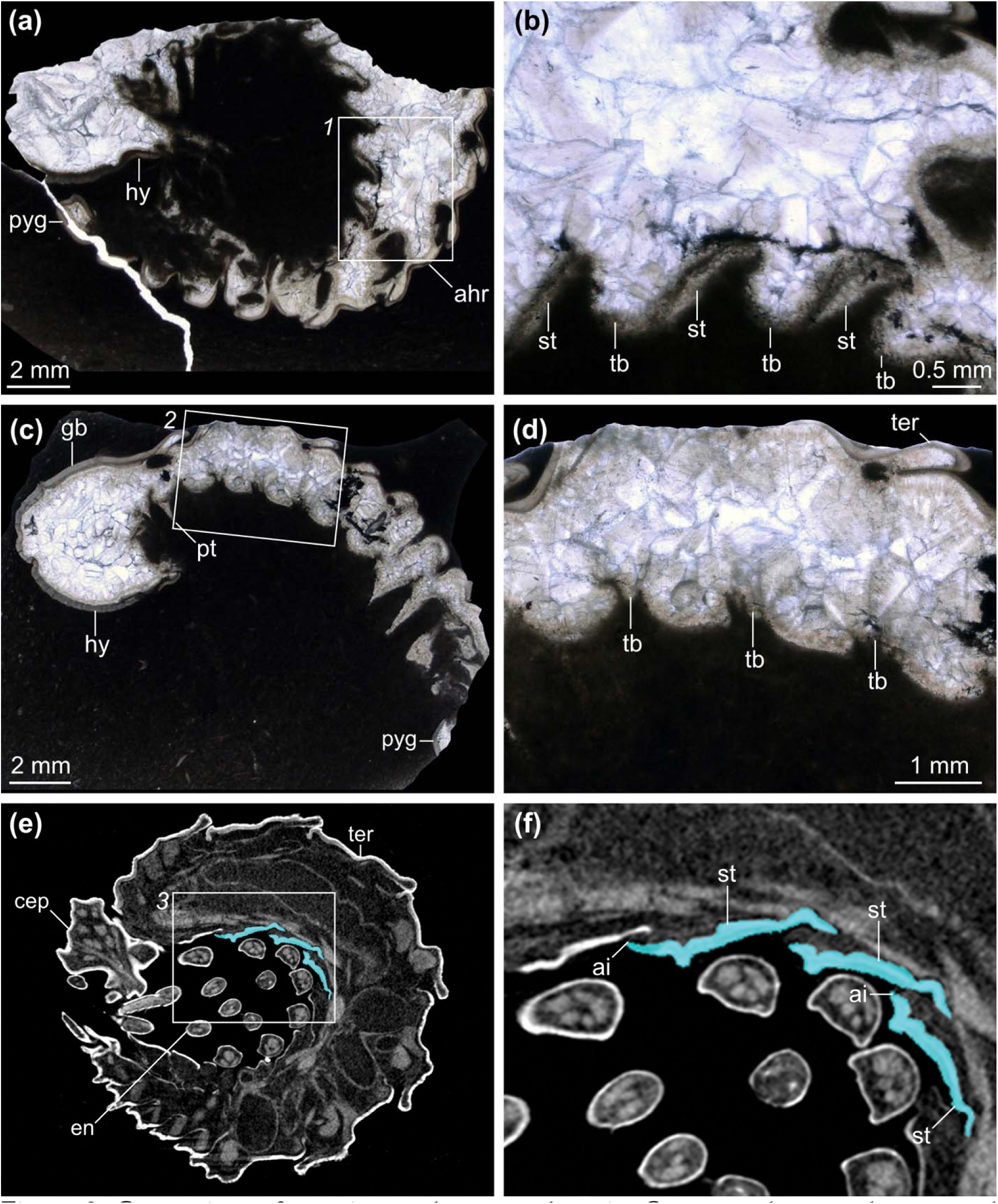
Comparison of appendage attachment of glomerid millipede and terrestrial isopod. (a – c) Micro-CT of fully enrolled glomerid millipede (MCZ:IZ:165554). (a) Three-dimensional volume rendering. (b) Tomographic slice showing medial appendage attachment. (c) Tomographic slice showing appendage orientation during enrollment. (d – f) Micro-CT of fully enrolled terrestrial isopod (MCZ:IZ:90105). (d) Three-dimensional volume rendering. (e) Tomographic slice showing lateral appendage attachment. (f) Tomographic slice showing appendage orientation during enrollment. Abbreviations: att, attachment of appendage to body wall; en, endopodite; pt, protopodite.

MCZ:IP:158251 features five imbricating and serially arranged ventral structures that dip anteriorly at a 50° angle relative to each other and nearly perpendicular to the dorsal exoskeleton (Fig. 2b). Unlike the dorsal articulating half rings in the same specimen, the ventral structures do not directly overlap each other (Fig. 2b) but are continuous between them based on the distribution of the fibrous calcite. We interpret these ventral features as direct evidence of sternites in *C. pleurexanthemus*, expressed as the thickened exoskeletal plates, connected on the anterior and posterior margins by transverse series of flexible tendinous bars (Fig. 2b). Another specimen of *C. pleurexanthemus* (MCZ:IP:158227) shows a similar sagittal section along the midline of the body, but here the trunk is only partially enrolled, a position that informs the position of the ventral structures under a different configuration (Fig. 2c). MCZ:IP:158227 also preserves five sets of repeating structures, but because of the more abaxial position of the thin section, as indicated by the protopodites seen posteriorly, the sternites are not visible. The anterior ventral structures are underneath the slightly enrolled first five tergites and consist of corrugated bulges with a shorter ventrally concave region between each (Fig. 2c, d). The lesser degree of trunk flexure in MCZ:IP:158227 shows that the sternites would be parallel relative to the dorsal exoskeleton in a fully prone position (Fig. 2c, d).

Comparisons with 3D datasets of partially and completely enrolled isopods and millipedes supports the interpretation of the ventral morphology of *Ceraurus pleurexanthemus* as all taxa display the same pattern of anterior imbrication of sternites during enrollment despite their different morphologies (Figs. 2, 3). The terrestrial isopod displays a more complex sternite morphology than trilobites, with a row of paired rectangular plates (Fig. 3a – c) rather than the single row of hourglass shaped sternites (Whittington, 1993; Ortega-Hernández and Brena, 2012). The anterior edge of the isopod sternite dips ventrally during enrollment below the posterior margin of the preceding plate (Figs. 2e, f, 3c). Glomerid millipedes have wish-bone-shaped sternites (Fig. 3d – f) with limbs emerging from protopodite/coxa cavities between adjacent ventral plates (Fig. 3f). The elongate lateral portions of each sternite align with the adjacent pleurites (Fig. 3e; Supplemental Fig. 1) and imbricate anteriorly during enrollment (Fig. 3f).

**Figure 3.**
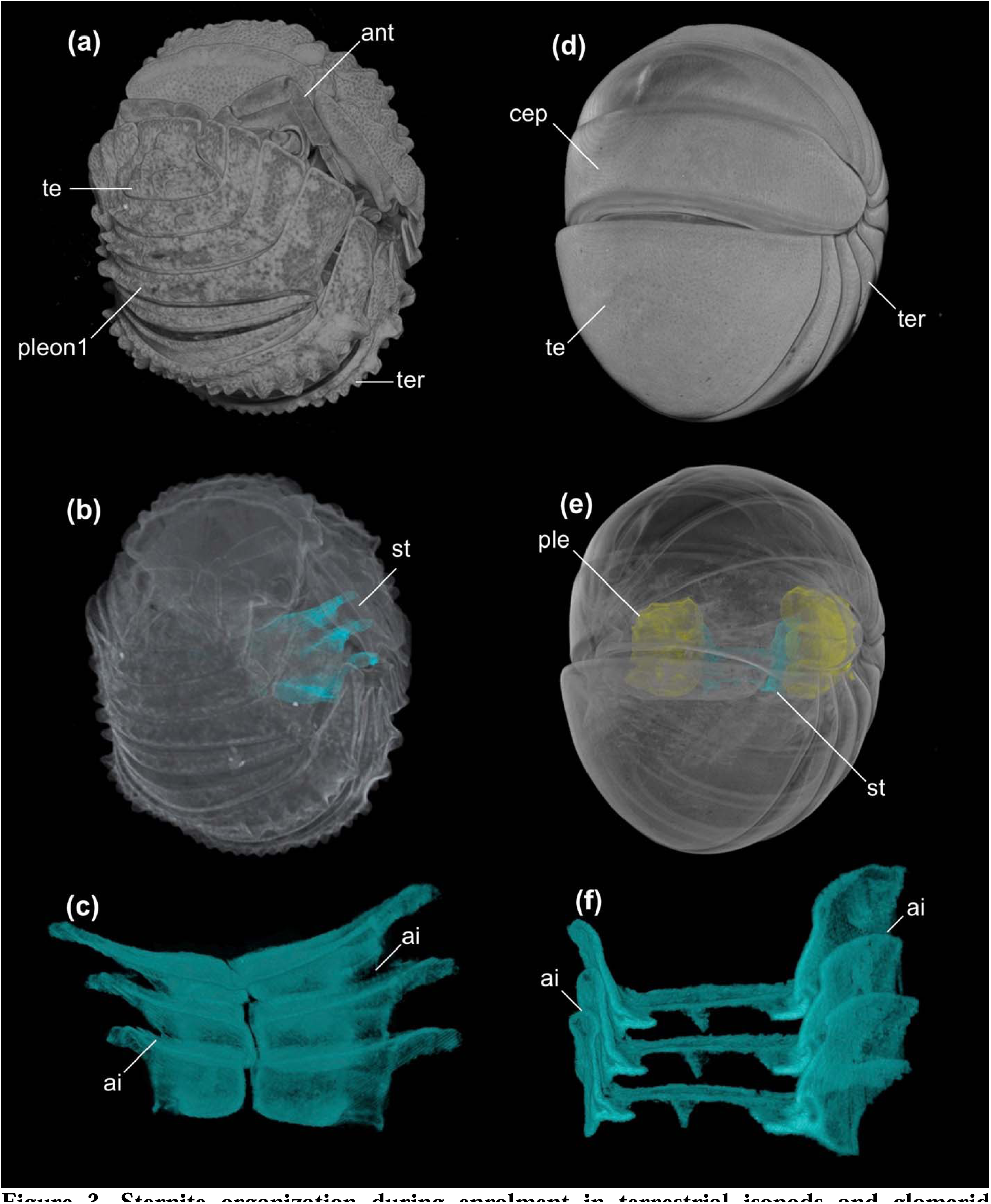
Sternite organization during enrolment in terrestrial isopods and glomerid millipedes. (a – d) Micro-CT scan of terrestrial isopod MCZ:IZ:90105. (a) Reconstruction of 90% enrolled specimen. (b) Micro-CT of full specimen with three segmented sternites (blue highlight). (c) Micro-CT segmented sternites from three segments. (d – f) Micro-CT scan of glomerid millipede MCZ:IZ: 165554. (d) Reconstruction of completely enrolled specimen. (e) Reconstruction of full specimen with three segmented sternites (blue highlight) and pleurites (yellow highlight). (f) Micro-CT segmented sternites from three segments. Abbreviations: ai, anterior imbrication; ant, antenna; cep, cephalon; ple, pleurite; st, sternites; te, telson; ter, tergite.

### Protopodite morphology and preservation

Exsagittal thin sections of Walcott-Rust trilobites such as MCZ:IP:158240 (*Ceraurus pleurexanthemus*; Fig. 4a, b) and MCZ:IP:104956 (*Flexicalymene senaria*; Fig. 4c, d) show the lateral ventral morphology in three-dimensions with exceptional detail. The hypostome and articulating half rings of the tergites are visible in these specimens, similarly to the sagittal thin section in MCZ:IP:158251 (Fig. 2a), but the posterior projections of the hypostome indicates that the more abaxial position near the lateral margin of the axial lobe (Fig. 4a, c). In both MCZ:IP:158240 and MCZ:IP:104956, a series of serially repeating wedge-shaped ventral structures are associated with each of the tergites (Figs. 4a, 5c, d). The presence of fibrous calcite defining these structures indicates that they were originally non-biomineralized (Losso et al., 2023). The wedge-shaped ventral structures are widest dorsally and taper ventrally to a point at a 40-50° angle (Fig. 4b, d). Specimen MCZ:IP:104956 of *F. senaria* displays a series of 22 wedge-shaped structures, four of which are associated with the cephalon, but none are visible beneath the pygidium (Fig. 4c). MCZ:IP:104956 is 65% enrolled and the series of wedge-shaped structures are angled slightly anteriorly (Fig. 4c). The anterior most wedges have a straight anterior margin that gently curves anteriorly at their mid-section (Fig. 4d). The posterior margin of the wedge is similarly curved, which allows for the succeeding wedges to fit snuggly against one another when in direct contact (Fig. 4d). The terminal tip of the wedge is slightly enlarged and bulbous (Fig. 4d).

**Figure 4.**
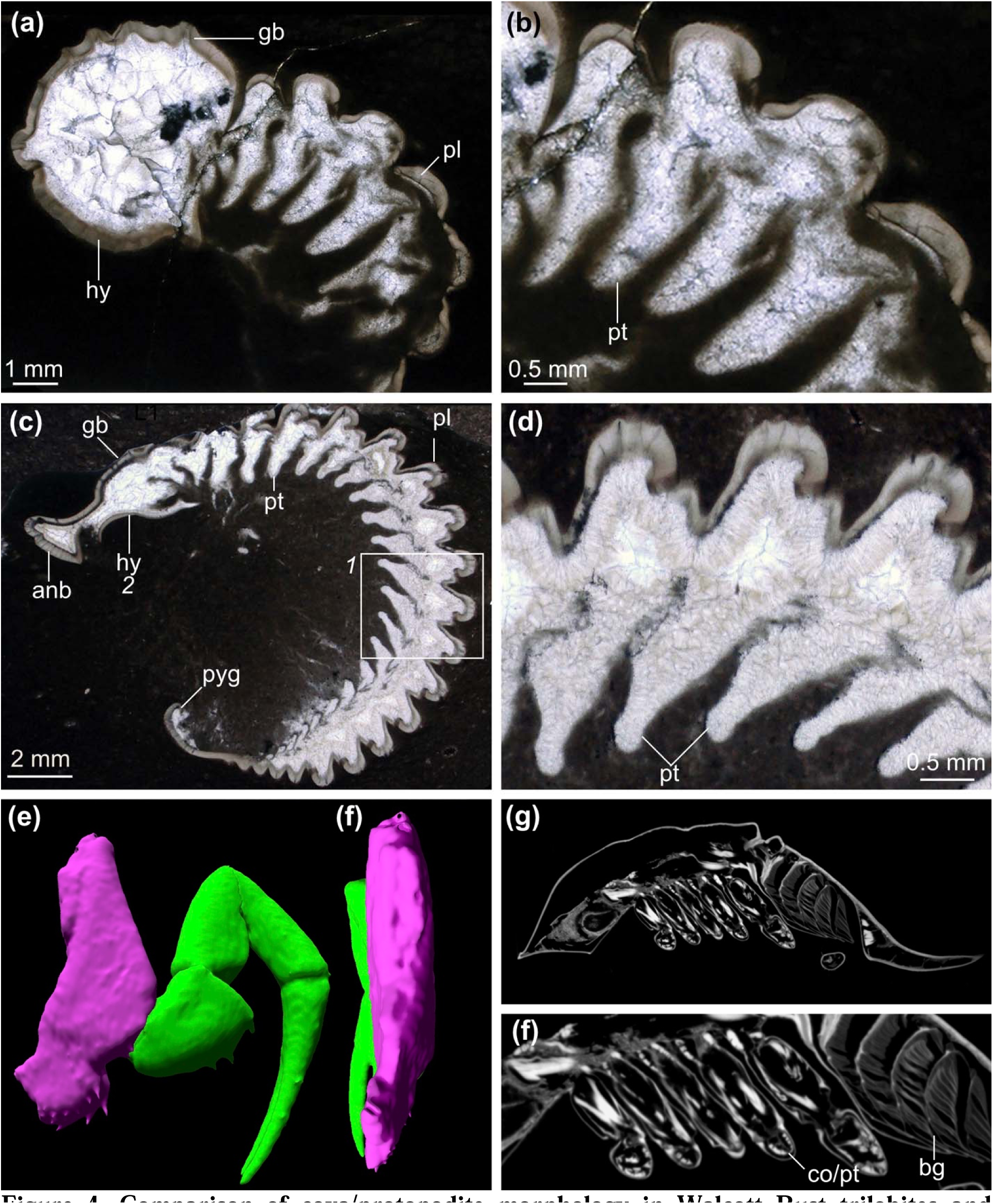
Comparison of coxa/protopodite morphology in Walcott Rust trilobites and *Limulus polyphemus* in lateral section. (a) Photomicrograph of *Ceraurus pleurexanthemus*, an exsagittal thin section of showing protopodites in cross section from a lateral view (MCZ:IP:158240). (b) Magnification of protopodites. (c) Photomicrograph of *Flexycalymene senaria*, a transverse thin section of showing protopodites in dorsal view (MCZ:IP:110918). (d) Magnification of protopodites in box 1 of (c). (e) Micro-CT segmentation of *Limulus polyphemus* (MCZ:IZ: 41112) showing anterior view of walking leg two including coxa/protopodite (purple highlight) and endopodite (green highlight). (f) Micro-CT segmentation of *Limulus polyphemus* (MCZ:IZ: 41112) showing medial view of walking leg two. (g) Tomographic slice of *Limulus polyphemus* (MCZ:IZ: 41112) in exsagittal view showing lateral section of protopodite. (h) Magnification of coxa/protopodite in lateral section. Abbreviations: anb, anterior band of cranidium; co, coxa; gb, glabellae; hy, hypostome; pl, pleural lobe; pt, protopodite; pyg, pygidium.

**Figure 5.**
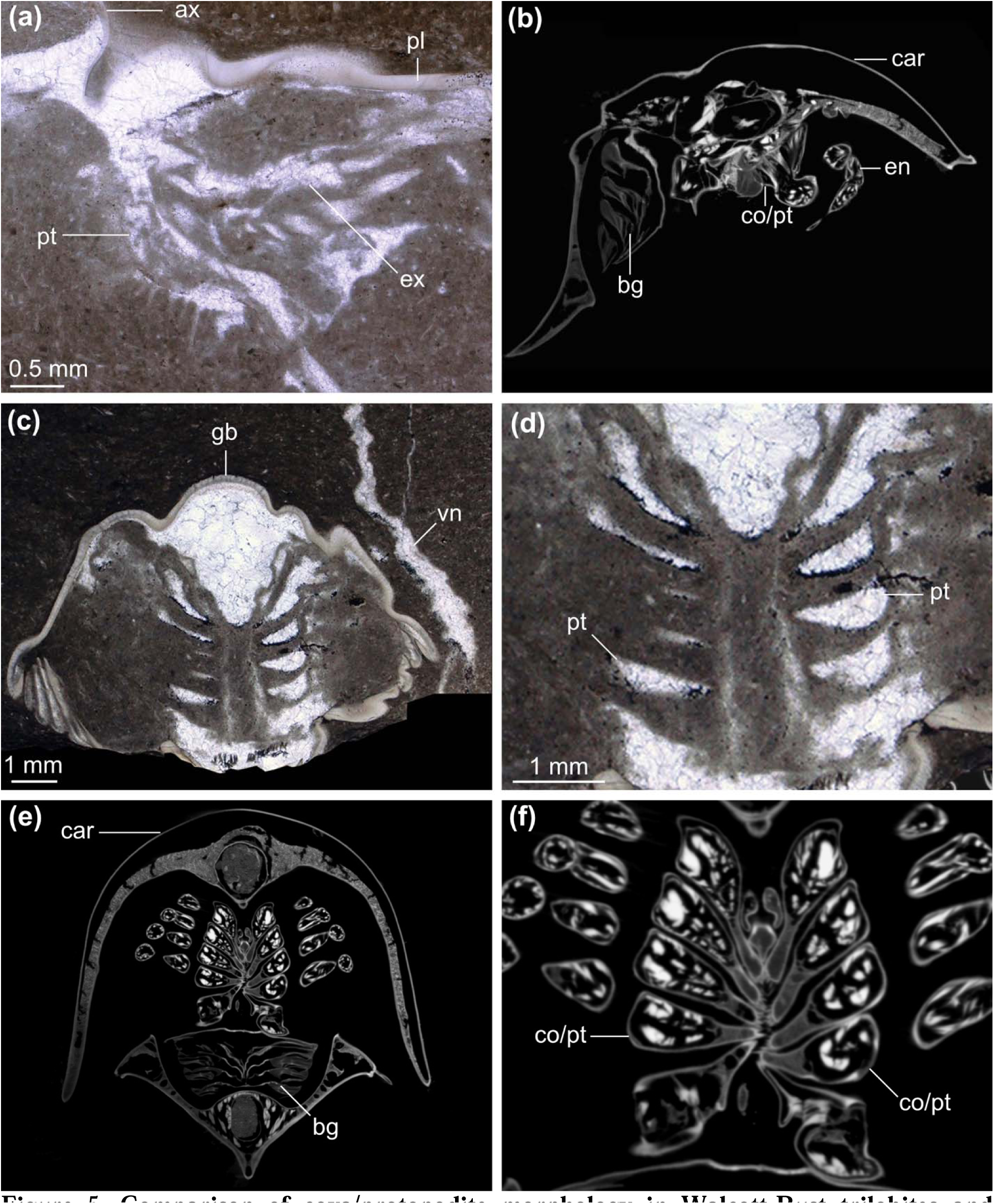
Comparison of coxa/protopodite morphology in Walcott-Rust trilobites and *Limulus polyphemus* in anterior and coronal sections. (a) Photomicrograph of *Ceraurus pleurexanthemus*, a transverse thin section of showing an anterior-posterior view of the protopodite (MCZ:IP:110933). (b) Tomographic slice of *Limulus polyphemus* (MCZ:IZ:41112) showing anterior view of coxa/protopodite. (c) Photomicrograph of *Flexicalymene senaria*, a transverse thin section showing coronal view of four protopodites (MCZ:IP:110918). (d) Magnification of protopodites of MCZ:IP:110918. (e) Tomographic slice of *Limulus polyphemus* (MCZ:IZ: 41112) showing coronal view of coxae/protopodites. (f), Magnification of coxae of MCZ:IZ:41112. Abbreviations: ax, axial lobe; car, carapace; co/pt, coxa/protopodite; bg, book gill; ex, exopodite; gb, glabella; pl, pleural lobe; pt, protopodite; vn, calcite vein.

Based on their taphonomy and morphology, we interpret the serially repeating wedge-shaped structures observed in both *C. pleurexanthemus* and *F. senaria* as direct evidence of three-dimensionally preserved protopodites, namely the part of the arthropodized biramous appendage that is in direct contact with the body wall (Boxshall, 2004), as observed in a cross-sectional view from an exsagittal plane. This interpretation is supported by the precise association of a wedge with each tergite (Fig. 4c) and the abaxial position relative to the axial lobe of the dorsal exoskeleton which can also be seen in a transverse thin section of *C. pleurexanthemus* showing an anterior view of the proximal portion of the appendage (Fig. 5a).

The repeating wedge-shaped structures are not part of the biomineralized dorsal exoskeleton, such as muscle attachment sites or apodemes (Whittington et al., 1997; Edgecombe and Sherwin, 2001; Siveter et al., 2021) as the original calcite would be clearly visible similar to the tergites and sternites (Fig. 2b, c). Comparisons with additional thin section specimens of *C. pleurexanthemus* further strengthen the interpretation of the wedges as protopodites. Specimen MCZ:IP:110933 shows a clear and unobstructed view of one biramous appendage (Fig. 5a), which shows that the laterally splayed protopodites of *C. pleurexanthemus* are subtriangular in anterior view with a nearly vertical medial margin and horizonal ventral margin. The medial margin is studded with gnathobasic spines, and the ventral edge is marked by elongate endites (Fig. 5a). The exopodite is visible dorsally and the endopodite extends from the distal margin of the protopodite (Fig. 5a). The protopodite of MCZ:IP:110933 extends from the lateral edge of the dorsal exoskeleton’s axial lobe and partially into the pleural lobe, which closely correlate with the position of the wedge-shaped structures seen in exsagittal thin sections (Fig. 4a, c). One specimen (MCZ:IP:110918) of *Flexicalymene senaria* sectioned in coronal view shows a series of wedge-shaped structures whose apex points towards the midline of the body (Fig. 5c, d). The position within the body and the comparison with the two other views (Fig. 4c, d), support the interpretation of these structures also represent the protopodites as observed from a dorsal section, which further confirms their wedge-shaped three-dimensional organization.

Comparisons with the 3D morphology of *Limulus polyphemus* supports the interpretation of the ventral structures seen in *Ceraurus pleurexanthemus* and *Flexicalymene senaria* as exsagittal and coronal views of protopodites based on the wedge-shaped morphology of the coxa/protopodite (Fig. 4e, f). The coxa/protopodite of the walking legs in *L. polyphemus* are dorsoventrally elongate with a large area of attachment to the body wall, the dorsal edge is broad cross section decreasing in width ventrally (Fig. 4e, f). In cross section from an exsagittal view, the coxae/protopodite are broadest dorsally, tapering ventrally (Fig. 4g, h). The coxae/protopodite are also wedge-shaped in coronal view, narrowest near the body wall and widening distally (Fig. 5e, f), a condition that is also seen in *Flexicalymene senaria* (Fig. 5c, d).

## Discussion

Walcott-Rust trilobites reveal new insights into the ventral morphology of trilobites, with direct implications for understanding their adaptations for enrollment. A single row of hour-glass shaped sternites are known throughout Trilobitomorpha such as *Arthroaspis bergstroemi* (Stein et al., 2013), *Misszhouia longicaudata* (Zhang et al., 2007; Mayers et al., 2019), and *Sinoburius lunaris* (Chen et al., 2019). The pyritized olenid trilobite *Triarthrus eatoni* preserves appendages, sternites and tendinous bars (Whittington and Almond, 1987), which closely resemble those found in the pliomerid *Placoparia cambriensis* (Whittington, 1993). These examples of preserved sternites are only observable in either ventral or dorsal view due to their preservation in compacted body fossils, therefore not providing information about the three-dimensional morphology, position within the body or movement during enrollment. The Walcott-Rust specimens provide complementary views of the sternites and protopodites that allow to reconstruct their three-dimensional overall morphology.

### Sternite position during enrollment

All known cases of sternite preservation in trilobites and non-biomineralized trilobitomorphs point towards the same broad pattern of morphological organization, in which the sternites are successively arranged in an axial row that runs parallel to the dorsal exoskeleton, and which are separated by flexible tendinous bars (Fig. 2c, d) (Whittington, 1993; Zhang et al., 2007; Stein et al., 2013; Mayers et al., 2019; Chen et al., 2019). The sternites are cuticular and have a thinner constitution than the dorsal exoskeleton; however, the ventral side of the body would not be able to physically accommodate the entire sternite series during complete enrollment while maintaining their outstretched disposition without producing excessive tension on either the ventral side, due to over compression, or the dorsal side, due to over extension (Fig. 5f). Instead, the ventral data from Walcott-Rust trilobites demonstrate that the sternites and tendinous bars become corrugated in the transition between prone position to partial and complete enrollment (Fig. 2a, b), with the anterior edge of the sternite angling ventrally and the flexible tendinous bar bulging (Fig. 6a-d). Critically, the same configuration between the sternites and arthrodial membranes are also observed in isopods (Fig. 3a – c) and millipedes (Fig. 3d – f), with the anterior edge of the sternites dipping ventrally to accommodate tight and complete enrollment. These comparisons indicate that despite the distant phylogenetic relationships between trilobites (extinct stem-group chelicerates), isopods (extant crustaceans) and millipedes (extant myriapods), these euarthropods share fundamentally similar exoskeletal ventral adaptations that facilitate protective enrollment. These findings evidence a striking case of convergent evolution that is heavily influenced by the mechanical requirements and limitations necessary to achieve complete encapsulations in euarthropods, which have been extensively investigated in terms of their dorsal exoskeletal morphology (e.g., Esteve et al., 2011, 2017, 2018; Chipman and Drage, 2023), but never from the perspective of the ventral anatomy until now. Repeated convergent evolution of the same mechanism in three distantly related euarthropods demonstrates the constraints of achieving complete enrollment with a rigid exoskeleton, and simultaneously the evolutionary advantages that this strategy must confer. Even more notably sternite morphology varies between the three taxa, from a single row of hourglass-shaped plates in trilobites (Fig. 2a), paired rectangular plates in isopods (Fig. 3c), to wish-bone-shaped sternites in glomerids (Fig. 3f), yet all species enroll utilizing the same basic mechanism.

**Figure 6.**
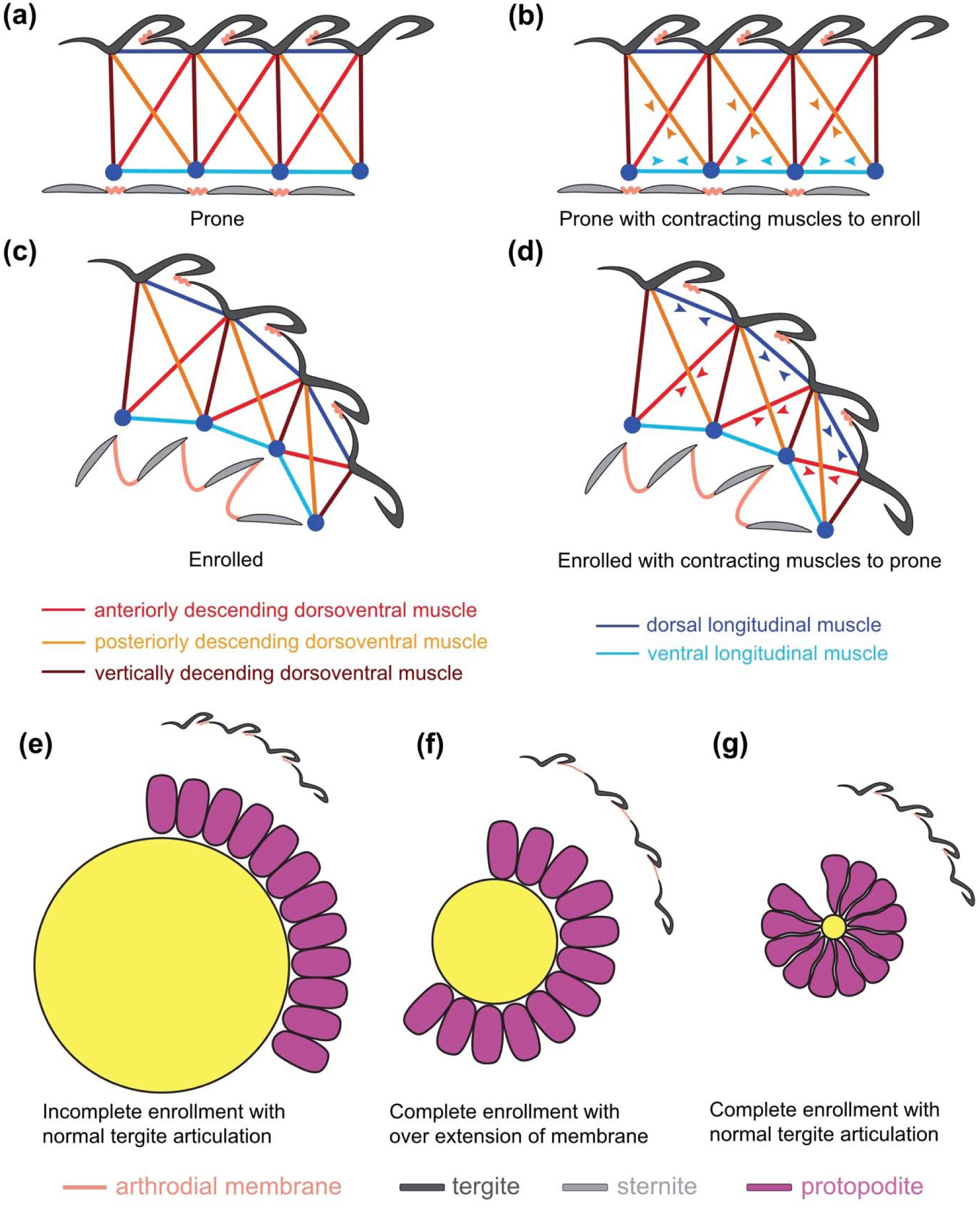
Functional morphology of trilobite enrollment. (a – d), Hypothesized muscle attachment in trilobites with contraction indicated by arrow heads. (a) Prone position. (b) Prone position showing contraction along the VLM and DVP to enroll. (c) Enrolled position with corrugation sternites dipping anteriorly and extended tenuous bars. (d) Enrolled position showing contraction of DLM and DVA to extend the body. (e – g) Diagram showing impact of protopodite morphology on complete and tight enrollment. (e) Oval protopodites restrict complete enrollment under normal tergite extension. (f) Oval protopodites with maximum contraction of the ventral edge cause excessive dorsal extension, exposing the arthrodial membrane. (g) Wedge-shaped protopodites based on MCZ:IP:104956 facilitate complete and tight enrollment without over extension of dorsal arthrodial membrane.

### Functional implications for trilobite musculature during enrollment

The new insights into the ventral exoskeletal organization of the trilobite exoskeleton have direct implications for understanding the functional morphology of the trunk musculature during enrollment (Fig. 6). Three proposed dorsal-ventral muscles would attach from tergites to sternites based on crustacean analogs; the anteriorly descending muscle, the vertically descending muscle and the posteriorly descending muscle (Cisne, 1981; Whittington, 1993). Longitudinal ventral muscles have been proposed to be on either side of the body of trilobites, attaching to the tendinous bar (Beecher, 1902; Størmer, 1939; Hupe, 1953; Whittington, 1993). The reconstruction of an exceptionally preserved specimen of *Placoparia cambriensis* with soft tissues in sagittal cross section (Whittington, 1993) bears striking similarities to MCZ:IP:158227 which is not fully enrolled (Fig. 2c). However, the fully enrolled reconstruction illustrates the ventral anatomy exactly the same as in the prone position (Whittington, 1993) despite the Walcott-Rust trilobites and MCZ:IP:158251 having been published nearly 100 years before with the corrugated ventral structures highlighted (Walcott 1881). A proposed mechanism of enrollment and extension for trilobites relied on the contraction of the longitudinal ventral muscle would bring the sternites closer together, enrolling of the body versus contraction of the dorsal longitudinal muscle that would extend the body to a prone position (Whittington, 1993) (Fig 6).

In extant euarthropods that completely enroll their body similarly to trilobites, such as glomeriid millipedes and terrestrial isopods, contraction of the longitudinal ventral muscle flex the body ventrally (Manton, 1961). However, the sternites are too long (sag.) to allow complete enrollment without substantial overlap between them (Fig. 3c, f). In this context, the sagittal thin section of *C. pleurexanthemus* (MCZ:IP: 158251) demonstrates that trilobite enrollment was accomplished by ventrally dipping the anterior margin of the sternites, producing a corrugated outline of sternites and arthrodial membrane (Fig. 3c, f). Based on their basic functional morphology, euarthropod enrollment is achieved through contraction of the longitudinal ventral muscle and the vertically descending muscle, bringing sternites closer together and raising the posterior margin dorsally (Fig. 6b, c). Returning to the prone position is accomplished through contraction of the longitudinal dorsal muscle and anteriorly descending muscle, which brings the tergites closer together and raises the anterior margin of the sternite to a horizontal position (Fig. 6d).

### Functional implications of wedge-shaped protopodites

The overall morphological organization of the trilobite protopodite has been elusive because of its non-biomineralized nature and proximity to the midline, with three-dimensional structure being especially difficult to reconstruct because of a lack of suitable trilobite fossils that clearly show this structure (Whittington, 1975; Zeng et al., 2017; Holmes et al., 2020; Bicknell et al., 2021; Hou et al., 2021; Siveter et al., 2021). This has left a gap in our understanding of locomotion, enrollment, and feeding autecology because of the crucial role of the protopodite (Bruton and Haas, 1999; Ortega-Hernández and Brena, 2012; Bicknell et al., 2018, 2021). Appendages are most frequently preserved in highly compressed anterior or posterior views such as *Olenoides serratus* from the Burgess Shale (Whittington, 1975; Hou et al., 2021), *Redlichia rex* (Holmes et al., 2020) from Emu Bay, or *Hongshiyanaspis yiliangensis* (Zeng et al., 2017). The lack of detailed information regarding the cross section of the trilobite protopodite has resulted a variety of hypothetical morphological interpretations. For instance, the cross section the trilobite protopodite has been reconstructed as having disparate shapes, including oval (e.g. Whittington, 1975; Whittington and Almond, 1987; Bruton and Haas, 1999; Ortega-Hernández and Brena, 2012; Bicknell et al., 2021; Schmidt et al., 2021), square (Hou et al., 2021), or this aspect has been omitted them altogether because of the lack of available data (Whittington, 1993).

Wedge-shaped protopodites may be common across Trilobita and even more broadly within euarthropods. Specimens of *Isotelus* (DeKay, 1824) are preserved with ventral views of three-dimensional protopodites (Mickleborough, 1883; Raymond, 1920) which appear as thin transverse to anterolateral bars, a similar condition to that seen in *Triarthrus eatoni* (Hall, 1838) (Whittington and Almond, 1987). Wedge-shaped protopodites can explain this view, as the *Isotelus* specimens show a coronal cross section similar to *Flexycalymene senaria* MCZ:IP:110918 (Fig. 4C). The thin and elongate appears of protopodites in *Isotelus* and *T. eatoni* is the result of the coronal view through the ventral edge or middle of the structure resulting in a superficially noodle like appearance. The Silurian trilobite *Dalmanites* sp. from the Herefordshire Biota is preserved as three-dimensional calcite casts visualized through serial sectioning (Siveter et al., 2021) similar to the preservation seen in the Walcott-Rust fossils (Losso et al., 2023). This allows segmentation of individual appendages which display wedge-shaped protopodites resembling those of *Flexicalymene senaria* and *Ceraurus pleurexanthemus* (Fig. 4a-d). Wedge-shaped protopodites are also found in other trilobitomorphs, such as xandarellids based on a specimen preserved as a three-dimensional external mold in displaying a coronal view of the appendages which plunge into the matrix (Ortega-Hernández et al., 2017). The protopodites appear elongate and slender, similar to *Isotelus* and *T. eatoni* discussed above. Given the similar orientation to those specimens, the wedge-shaped protopodite morphology can also account for this specimen with only a 2D view of the structures being visible from the surface. An exception to this appears to be *Agnostus pisiformis* (Linnaeus, 1747) which has an oval coronal cross section (Müller and Walossek, 1987), but determining protopodite cross section from the published literature is difficult because the required views are rarely illustrated for extant species and rarely preserved in fossil specimens.

The widely reproduced oval cross section for the protopodite (Whittington, 1975; Whittington and Almond, 1987; Bruton and Haas, 1999; Ortega-Hernández and Brena, 2012; Bicknell et al., 2021; Schmidt et al., 2022) would severely hamper the ability of trilobites enroll effectively. The oval protopodite would make it impossible for trilobites to achieve complete enrollment under the normal observed range of motion of the tergites (Fig. 6E), or alternatively, would require the dorsal side of the body to overextend significantly, leaving open gaps between the articulating tergites and exposing the delicate arthrodial membrane to predators (Fig. 6F). In this context, the distinctively wedge-shaped protopodites of *Flexicalymene senaria* and *Ceraurus pleurexanthemus* (Fig. 3A – D) would play a critical role during enrollment by facilitating a tight body flexure, but without causing dorsal over extension thanks to their narrow ventral margin and form-fitting shape relative to each other (Fig. 6G). Comparisons with the three-dimensional appendage morphology of glomerid millipedes and terrestrial isopods indicate that these extant taxa do not have a wedge-shaped protopodite but differ from trilobites in having medially-(Supplemental Figs. 1, 2a – c) or laterally- (Supplemental Fig. 2d – f) attached appendages as opposed to intermediate condition as in trilobites (Fig. 4a) and other extinct trilobitomorphs. A critical difference between these extant taxa and trilobites, however, is the fact that the former do not utilize the trunk appendages for food processing, but instead employ the modified mandibles as mouthparts (Köiuhler and Alberti, 1990), whereas the entire limb series of trilobites has an active role in feeding based on the presence of well-developed gnathobasic spines along the food groove (Hegna, 2009; Bicknell et al., 2021). By contrast, the three-dimensional morphology of the trilobite protopodite is more similar to that of *Limulus polyphemus* both in terms of its transverse and dorsal sections (Fig. 3E-H; Fig. 4F, G), which does engage in aquatic gnathobasic feeding with the entire prosomal limb series (Bicknell et al., 2018, 2021). Based on these comparisons, we propose that the wedge shape of the trilobite protopodite reflects a unique functional tradeoff between the physical constrains required for tightly accommodating the appendages during complete enrollment (Fig. 6G), coupled with their pivotal role in food processing during gnathobasic feeding through the ventral groove of the body.

The ability of trilobites to enroll for protection represents an iconic adaptation that heavily influenced the long and successful evolutionary history of these extinct euarthropods. Our new data from the Walcott-Rust Quarry reveal for the first time that, in addition to the coaptative devices on the dorsal calcitic exoskeleton, trilobites also featured ventral morphological adaptations of the non-biomineralized sternites and biramous appendages that played a critical and multifaceted role in their mode of life. We find direct evidence that the fundamental mechanism of sternite corrugation that facilitate complete enrolment in trilobites is also expressed in extant glomerid millipedes and terrestrial isopods, showing a striking case of convergent evolution in phylogenetically distant euarthropod clades separated by hundreds of millions of years.

## Acknowledgements

We thank Adam Baldinger and Jessica Cundiff (Museum of Comparative Zoology, Cambridge, USA) for facilitating access to specimens.

## Supplemental Figures

**Figure 1.**
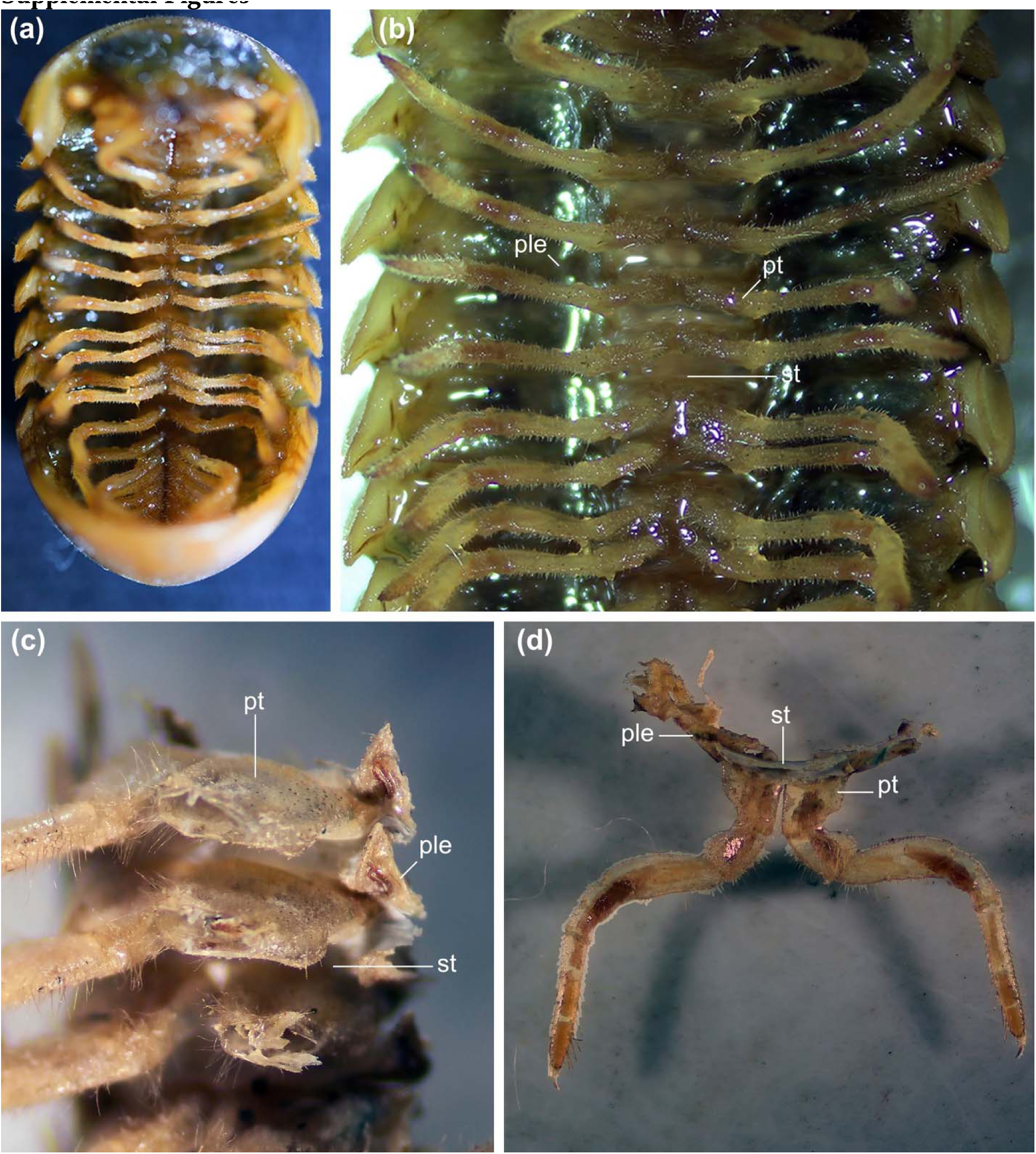
Protective enrollment in trilobites and extant euarthropods. (a) *Calymene fayettens* (MCZ:IP:5112). (b) *Isotelus maximus* (MCZ:IP:58). (c) *Phacops foecundus* (MCZ:IP:201074). (d) *Proetus bohemicus* (MCZ:IP:5264). (e) Terrestrial isopod (MCZ:IZ:90105). (f) Glomerid millipede (MCZ:IZ:165554).

**Figure 2.**
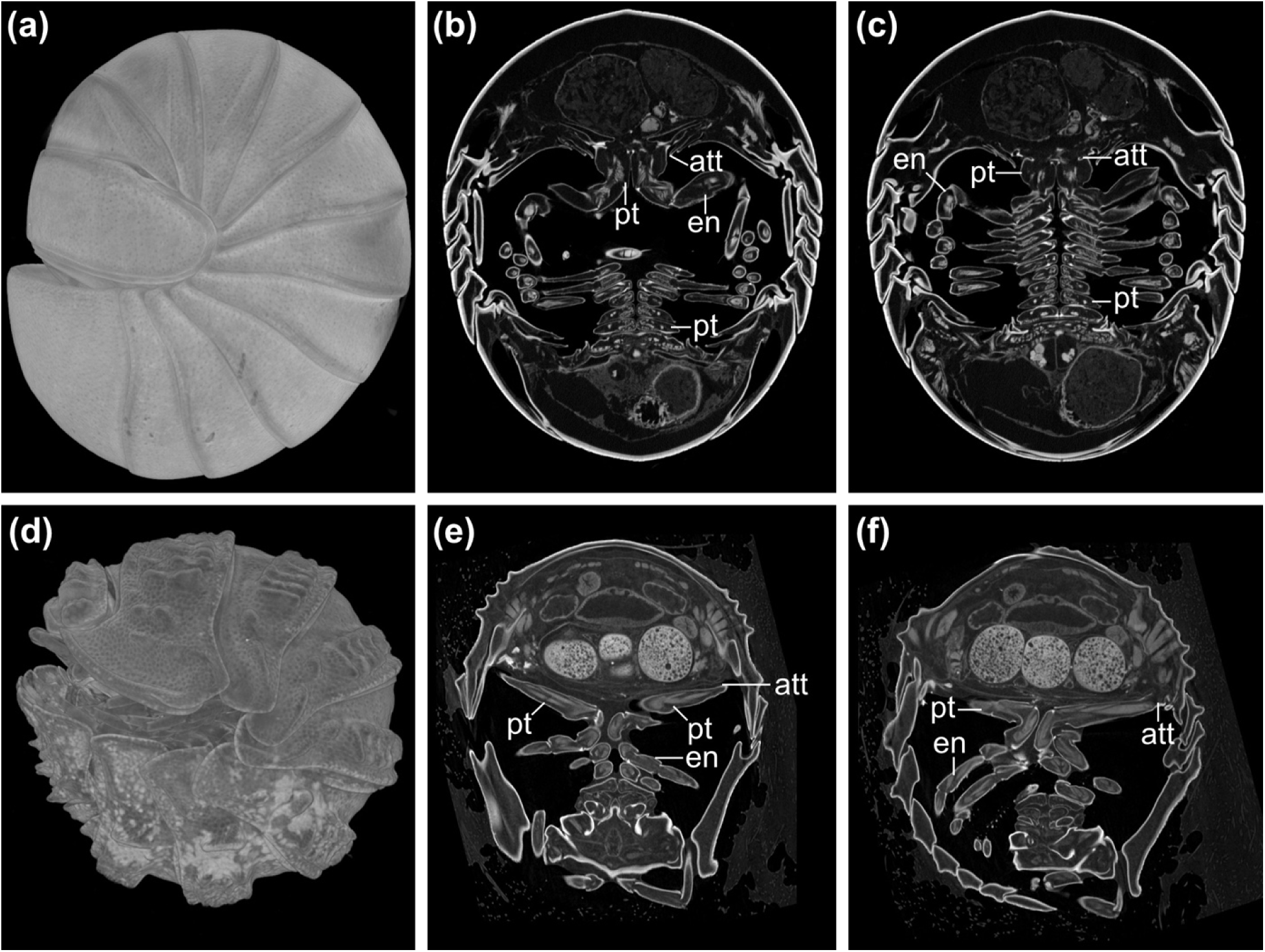
Comparison of sternites and tenuous bars in *Ceraurus pleurexanthemus* and terrestrial isopod in lateral exsagittal section. (a) Photomicrograph of MCZ:IP:158251, a sagittal thin section of a nearly completely enrolled specimen with preserved sternites and tendinous bars. (b) Photomicrograph of MCZ:IP:158251 showing magnification of sternites box 1 of (a). (c) Photomicrograph of MCZ:IP:158227, a sagittal thin section showing tendinous bars in partial enrollment. (d) Photomicrograph of MCZ:IP:158227 showing magnification of tendinous bars box 2 of (c). (e) Tomographic slice of isopod MCZ:IZ:90105 showing corrugation of sternites (blue highlight). (f) Magnification of sternites (MCZ:IP:90105). Abbreviations: ahr, articulating half ring; cep, cephalon; en, endopodite; gb, glabella; hy, hypostome; pyg, pygdium; st, sternite; tb, tenuous bar.

## References

1. Ballerio, A., and Grebennikov, V., 2016, Rolling into a ball: phylogeny of the Ceratocanthinae (Coleoptera: Hybosoridae) inferred from adult morphology and origin of a unique body enrollment coaptation in terrestrial arthropods: Arthropod Systematics & Phylogeny, v. 74, p. 23–52, doi:10.3897/asp.74.e31837.

2. Beecher, C.E., 1902, The ventral integument of trilobites: American Journal of Science, v. s4-13, p. 165–174, doi:10.2475/ajs.s4-13.75.165.

3. Bicknell, R.D.C., Holmes, J.D., Edgecombe, G.D., Losso, S.R., Ortega-Hernández, J., Wroe, S., and Paterson, J.R., 2021, Biomechanical analyses of Cambrian euarthropod limbs reveal their effectiveness in mastication and durophagy: Proceedings of the Royal Society B: Biological Sciences, v. 288, p. 20202075.

4. Bicknell, R.D.C., Ledogar, J.A., Wroe, S., Gutzler, B.C., Watson, W.H., and Paterson, J.R., 2018, Computational biomechanical analyses demonstrate similar shell-crushing abilities in modern and ancient arthropods: Proceedings of the Royal Society B: Biological Sciences, v. 285, p. 20181935, doi:10.1098/rspb.2018.1935.

5. Boxshall, G.A., 2004, The evolution of arthropod limbs: Biological Reviews, v. 79, p. 253–300, doi:10.1017/S1464793103006274.

6. Brökeland, W., Wägele, J.-W., and Bruce, N.L., 2001, Paravireia holdichi n. sp., an enigmatic isopod crustacean from the Canary Islands with affinities to species from New Zealand: Org. Divers. Evol.,.

7. Bruton, D.L., and Haas, W., 1999, The anatomy and functional morphology of Phacops (Trilobita) from the Hunsrück Slate (Devonian): Palaeontographica Abteilung A, v. 253, p. 29–75, doi:10.1127/pala/253/1999/29.

8. Chen, X., Ortega-Hernández, J., Wolfe, J.M., Zhai, D., Hou, X., Chen, A., Mai, H., and Liu, Y., 2019, The appendicular morphology of Sinoburius lunaris and the evolution of the artiopodan clade Xandarellida (Euarthropoda, early Cambrian) from South China: BMC Evolutionary Biology, v. 19, p. 165, doi:10.1186/s12862-019-1491-3.

9. Chipman, A.D., and Drage, H.B., 2023, Trilobites in rock enrol: a comment on ‘Developmental and functional controls on enrolment in an ancient, extinct arthropod’ by Esteve and Hughes (2023): Proceedings of the Royal Society B: Biological Sciences, v. 290, p. 20231547, doi:10.1098/rspb.2023.1547.

10. Cisne, J.L., 1981, Triarthrus eatoni (Trilobita): Anatomy of its exoskeletal, skeletomuscular, and digestive systems: Palaeontographica Americana, v. 9, p. 99–142.

11. DeKay, J.E., 1824, Oberservations on the structure of trilobites, and description of an apparently new genus. With notes on the geology of Trenton Falls: Annals of The Lyceum of Natural History of New York, v. 1, p. 174–189.

12. Edgecombe, G.D., and Sherwin, L., 2001, Early Silurian (Llandovery) trilobites from the Cotton Formation, near Forbes, New South Wales, Australia: Alcheringa: An Australasian Journal of Palaeontology, v. 25, p. 87–105, doi:10.1080/03115510108619215.

13. Esteve, J., GutiérrezLMarco, J.C., Rubio, P., and Rábano, I., 2018, Evolution of trilobite enrolment during the Great Ordovician Biodiversification Event: insights from kinematic modelling: Lethaia, v. 51, p. 207–217, doi:10.1111/let.12242.

14. Esteve, J., Hughes, N.C., and Zamora, S., 2011, Purujosa trilobite assemblage and the evolution of trilobite enrollment: Geology, v. 39, p. 575–578, doi:10.1130/G31985.1.

15. Esteve, J., Hughes, N.C., and Zamora, S., 2013, Thoracic structure and enrolment style in middle Cambrian Eccaparadoxides pradoanus presages caudalization of the derived trilobite trunk: Palaeontology, v. 56, p. 589–601, doi:10.1111/pala.12004.

16. Esteve, J., Rubio, P., Zamora, S., and Rahman, I.A., 2017, Modelling enrolment in Cambrian trilobites: Palaeontology, v. 60, p. 423–432, doi:10.1111/pala.12294.

17. Hall, J., 1838, Descriptions of two species of trilobites belonging to the genus Paradoxides: Journal of Science, v. 33, p. 199–202.

18. Hannibal, J.T., and Feldmann, R.M., 1981, Systematics and Functional Morphology of Oniscomorph Millipedes (Arthropoda: Diplopoda) from the Carboniferous of North America: Journal of Paleontology, v. 55, p. 730–746.

19. Hegna, T.A., 2009, The function of forks: Isotelus-type hypostomes and trilobite feeding: Lethaia, doi:10.1111/j.1502-3931.2009.00204.x.

20. Holmes, J.D., Paterson, J.R., and García-Bellido, D.C., 2020, The trilobite Redlichia from the lower Cambrian Emu Bay Shale Konservat-Lagerstätte of South Australia: systematics, ontogeny and soft-part anatomy: Journal of Systematic Palaeontology, v. 18, p. 295–334, doi:10.1080/14772019.2019.1605411.

21. Hou, X., Clarkson, E.N.K., Yang, J., Zhang, X., Wu, G., and Yuan, Z., 2008, Appendages of early Cambrian Eoredlichia (Trilobita) from the Chengjiang biota, Yunnan, China: Earth and Environmental Science Transactions of the Royal Society of Edinburgh, v. 99, p. 213–223, doi:10.1017/S1755691009008093.

22. Hou, J., Hughes, N.C., and Hopkins, M.J., 2021, The trilobite upper limb branch is a well-developed gill: Science Advances, v. 7, p. 1–8, doi:10.1126/sciadv.abe7377.

23. Hupe, P., 1953, Classe de Trilobites, in Traité de Paléontologie, Paris, Masson, v. III, p. 44–246.

24. Hyžný, M., and Dávid, A., 2017, A remarkably well-preserved terrestrial isopod (Peracarida: Isopoda: Armadillidiidae) from the upper Oligocene of Hungary, with remarks on the oniscidean taphonomy: Palaeontologia Electronica, doi:10.26879/615.

25. Köiuhler, H.-R., and Alberti, G., 1990, Morphology of the mandibles in the millipedes (Diplopoda, Arthropoda): Zoologica Scripta, v. 19, p. 195–202, doi:10.1111/j.1463-6409.1990.tb00255.x.

26. Linnaeus, C., 1747, Wästgöta-Resa, på Riksen höglofige ständers befallning förrättad år 1746. Med anmärkningar uti oeconmien, naturkunnogheten, antiquieter, inwånares seder och lefnads-sätt.: Stockholm, Lars Salvius.

27. Losso, S.R., and Ortega-Hernández, J., 2022, Claspers in the mid-Cambrian Olenoides serratus indicate horseshoe crab–like mating in trilobites: Geology, v. 50, p. 897–901, doi:10.1130/G49872.1.

28. Losso, S.R., Thines, J.E., and Ortega-Hernández, J., 2023, Taphonomy of non-biomineralized trilobite tissues preserved as calcite casts from the Ordovician Walcott-Rust Quarry, USA: Communications Earth & Environment, v. 4, doi:10.1038/s43247-023-00981-5.

29. Manton, S.M., 1961, The evolution of arthropodan locomotory mechanisms. Part 7. Functional requirements and body design in Colobognatha (Diplopoda), together with a comparative account of the diplopod burrowing techniques, trunk musculature and segmentation: Journal of the Linnean Society of London, Zoology, v. 44, p. 383–462, doi:10.1111/j.1096-3642.1961.tb01622.x.

30. Mayers, B., Aria, C., and Caron, J., 2019, Three new naraoiid species from the Burgess Shale, with a morphometric and phylogenetic reinvestigation of Naraoiidae (X. Zhang, Ed.): Palaeontology, v. 62, p. 19–50, doi:10.1111/pala.12383.

31. Mickleborough, J., 1883, Locomotory appendages of trilobites: Cincinnati Society of Natural History, p. 200–206.

32. Müller, K.J., and Walossek, D., 1987, Morphology, ontogeny, and life habit of Agnostus pisiformis from the Upper Cambrian of Sweden: Oslo, Universitetsforlaget, Fossils and strata no. 19, 124 p.

33. Ortega-Hernández, J., Azizi, A., Hearing, T.W., Harvey, T.H.P., Edgecombe, G.D., Hafid, A., and El Hariri, K., 2017, A xandarellid artiopodan from Morocco – a middle Cambrian link between soft-bodied euarthropod communities in North Africa and South China: Scientific Reports, v. 7, doi:10.1038/srep42616.

34. Ortega-Hernández, J., and Brena, C., 2012, Ancestral patterning of tergite formation in a centipede suggests derived mode of trunk segmentation in trilobites (P. K. Dearden, Ed.): PLoS ONE, v. 7, p. e52623, doi:10.1371/journal.pone.0052623.

35. Ortega-Hernández, J., Esteve, J., and Butterfield, N.J., 2013, Humble origins for a successful strategy: complete enrolment in early Cambrian olenellid trilobites: Biology Letters, v. 9, p. 20130679–20130679, doi:10.1098/rsbl.2013.0679.

36. Ramsköld, L., and Edgecombe, G.D., 1996, Trilobite appendage structure — Eoredlichia reconsidered: Alcheringa: An Australasian Journal of Palaeontology, v. 20, p. 269–276, doi:10.1080/03115519608619471.

37. Raymond, P.E., 1920, The Appendages, Anatomy, and Relationships of Trilobites: Memoirs of the Connecticut Academy of Arts and Sciences, v. 7.

38. Schmidt, M., Hou, X., Zhai, D., Mai, H., Belojević, J., Chen, X., Melzer, R.R., Ortega-Hernández, J., and Liu, Y., 2022, Before trilobite legs: Pygmaclypeatus daziensis reconsidered and the ancestral appendicular organization of Cambrian artiopods: Philosophical Transactions of the Royal Society B: Biological Sciences, v. 377, p. 20210030, 10.1098/rstb.2021.0030.

39. Schmidt, M., Hou, X., Zhai, D., Mai, H., Belojević, J., Chen, X., Melzer, R.R., Ortega-Hernández, J., and Liu, Y., 2021, Before trilobite legs: *Pygmaclypeatus daziensis* reconsidered and the ancestral appendicular organization of Cambrian artiopods: preprint, doi:10.1101/2021.08.18.456779.

40. Shear, W., Jones, T., and Wesener, T., 2011, Glomerin and homoglomerin from the North American pill millipede Onomeris sinuata (Loomis, 1943) (Diplopoda, Pentazonia, Glomeridae): International Journal of Myriapodology, v. 4, p. 1–10, doi:10.3897/ijm.4.1105.

41. Siveter, D.J., Fortey, R.A., Briggs, D.E.G., Siveter, D.J., and Sutton, M.D., 2021, The first Silurian trilobite with three dimensionally preserved soft parts reveals novel appendage morphology (X. Zhang, Ed.): Papers in Palaeontology, p. spp2.1401, doi:10.1002/spp2.1401.

42. Stein, M., Budd, G.E., Peel, J.S., and Harper, D.A., 2013, *Arthroaspis* n. gen., a common element of the Sirius Passet Lagerstätte (Cambrian, North Greenland), sheds light on trilobite ancestry: BMC Evolutionary Biology, v. 13, p. 99, doi:10.1186/1471-2148-13-99.

43. Størmer, L., 1939, Studies on trilobite morphology. Part I: The thoracic appendages and their phylogenetic significance: Norsk Geologisk Tidsskrift, v. 19, p. 143–274.

44. Suárez, M.G., and Esteve, J., 2021, Morphological diversity and disparity in trilobite cephala and the evolution of trilobite enrolment throughout the Palaeozoic: Lethaia, v. 54, p. 752– 761, doi:10.1111/let.12437.

45. Superina, M., and Loughry, W.J., 2012, Life on the Half-Shell: Consequences of a Carapace in the Evolution of Armadillos (Xenarthra: Cingulata): Journal of Mammalian Evolution, v. 19, p. 217–224, doi:10.1007/s10914-011-9166-x.

46. Walcott, C.D., 1881, The trilobite: new and old evidence relating to its organization: Bulletin of the Museum of Comparative Zoology at Harvard College, v. 8, p. 190–242.

47. Whittington, H.B., 1993, Anatomy of the Ordovician trilobite Placoparia: Philosophical Transactions of the Royal Society of London. Series B: Biological Sciences, v. 339, p. 109–118, doi:10.1098/rstb.1993.0008.

48. Whittington, H.B., 1975, Trilobites with appendages from the middle Cambrian, Burgess Shale, British Columbia: Fossils and Strata, p. 97–135.

49. Whittington, H.B., and Almond, J.E., 1987, Appendages and Habits of the Upper Ordovician Trilobite Triarthrus eatoni: Philosophical Transitions of the Royal Society of London, v. 317, p. 1–46.

50. Whittington, H.B., Moore, R.C., Geological Society of America, and Paleontological Society (Eds.), 1997, Treatise on Invertebrate Paleontology O: Lawrence, Kans, Univ. of Kansas Pr, Treatise on invertebrate paleontology Arthropoda 1. Trilobita Founder: Raymond C. Moore. Prepared under the sponsorship of The Geological Society of America, Inc., The Paleontological Society … ; Pt. O Vol. 1, 530 p.

51. Yochelson, E.L., 1998, Charles Doolittle Walcott, Paleontologist: Kent, Ohio, The Kent State University Press.

52. Zeng, H., Zhao, F., Yin, Z., and Zhu, M., 2017, Appendages of an early Cambrian metadoxidid trilobite from Yunnan, SW China support mandibulate affinities of trilobites and artiopods: Geological Magazine, v. 154, p. 1306–1328, doi:10.1017/S0016756817000279.

53. Zhang, X.-L., Shu, D.-G., and Erwin, D.H., 2007, Cambrian naraoiids (Arthropoda): morphology, ontogeny, systematics, and evolutionary relationships: Journal of Paleontology, v. 81, p. 1–52, doi:10.1666/06-082.1.

